# Evaluating Differences in Elastic Modulus of Regenerated and Uninjured Mouse Digit Bone through microCT Density-Elasticity Calculation and Nanoindentation Testing

**DOI:** 10.1101/2021.07.26.453818

**Authors:** Kevin F Hoffseth, Emily Busse, Michelle Lacey, Mimi C Sammarco

**Author notes:** Co-Corresponding: Kevin F. Hoffseth, Mimi Sammarco.

## Abstract

Bone is an essential, healing structure in vertebrates that ensures daily function. However, the regenerative capacity of bone declines with age, compromising quality of life in the elderly and increasing cost of care. Here, for the first time, the elasticity of regenerated bone in a mouse digit amputation model is evaluated in order to better investigate biomechanics of skeletal regeneration. Amputation of the distal one third of the digit (third phalangeal element – P3) results in de novo regeneration of the digit, where analyzing the structural quality of this regenerated bone is a challenging task due to its small scale and triangular shape. To date, the evaluation of structural quality of the P3 bone has primarily focused on mineral density and bone architecture. This work describes an image-processing based method for assessment of elasticity in the whole P3 bone by using microcomputed tomography-generated mineral density data to calculate spatially discrete elastic modulus values across the entire P3 bone volume. Further, we validate this method through comparison to nanoindentation-measured values for elastic modulus. Application to a set of regenerated and unamputated digits shows that regenerated bone has a lower elastic modulus compared to the uninjured digit, with a similar trend for experimental hardness values. This method will be impactful in predicting and evaluating the regenerative outcomes of potential treatments and heightens the utility of the P3 regenerative model.

## 1. Introduction

A crucial focus in the investigation of bone tissue regeneration and healing is that of bone quality, and, in particular, how “good” are the structural properties of regenerated bone tissue. Assessment of bone structural quality remains a critical challenge in research targeting the problems associated with bone health and aging, which often negatively affects quality of life in cascading fashion. In 2010, an estimated 10 million adults over the age of 50 suffered from osteoporosis (Wade et al., 2014).Patients with osteoporosis suffer from increasing bone loss and function, followed by an increased chance of fracture during a fall, accounting for approximately 1.5 million fractures annually in the USA (Riggs and Melton, 1995). The medical cost of osteoporosis and related fractures was estimated to be $22 billion dollars in 2008 (Blume and Curtis, 2011), with costs expected to increase along with the aging of the US population. A wide array of research seeks to improve the natural ability of bone tissue to repair and heal bone to prevent this decline in bone quality. This underscores the need for better methods to quantify bone quality before and after treatment, and for early diagnosis of bone diseases.

This research focuses on bone structural quality in an established skeletal regeneration model: amputation of the distal mouse digit tip resulting in patterned direct bone regeneration. In this model, the distal half of the third phalangeal element (P3) is amputated. Following amputation, the digit regenerates bone via direct ossification, whereas more proximal amputations fail to produce a regenerative response. The mouse digit amputation model has been a valuable model in studying mechanisms underlying bone regeneration and healing (Borgens, 1982; Brockes and Kumar, 2005; Bryant et al., 2002; Douglas, 1972; Fernando et al., 2011; Han et al., 2008; Illingworth, 1974; Said et al., 2004; Simkin et al., 2015; Singer et al., 1987). While work using this model has illuminated critical cell signaling factors that influence regenerative capacity on a molecular level, a gap remains in characterizing the biomechanical properties and functional structure of the regenerated bone.

Employment of micro-CT (µCT) imaging to investigate bone morphology has grown, coupled with image analysis and processing (Bouxsein et al., 2010). Methods of dual thresholding, 3D adaptive thresholding, genetic algorithms (Buie et al., 2007; Janc et al., 2013; Zhang et al., 2010), and shape recognition using deep learning techniques (Ambellan et al., 2019; Spampinato et al., 2017) have been used for segmentation of regions of interest. Regenerative outcomes in the mouse digit regeneration model have been largely quantified using µCT techniques coupled with ImageJ (Rasband, 1997), focusing on parameters such as length changes (Dawson et al., 2017), trabecular spacing, trabecular thickness, and volumetric change (Busse et al., 2019; Fernando et al., 2011; Sammarco et al., 2015; Sammarco et al., 2014; Simkin et al., 2017). Beyond micro-CT imaging, approaches such as DXA, CT, and MRI are also widely used for assessing bone mineral density and aspects of morphology. These approaches have enabled valuable insight into understanding how the regrown bone is formed under the regenerative process, however they lack solid links to mechanical properties and performance.

Comparison of calibrated volumetric bone mineral density (vBMD) to mechanical testing measurements can provide data that better quantifies and predicts the mechanical properties of bone (Figure 1). Relationships linking BMD to bone mechanical properties are critical for the development of physiologically accurate finite element models based on CT data and have been developed for use in the quantitative computed tomography to finite element pipeline (Knowles et al., 2016). Along these lines, density-modulus relationships have also been developed with various mechanical test methods and sample geometry, where continuous functions and power relationships return best fit with experimental results (Helgason et al., 2008). A wide selection of relationships has been developed for various types of bone tissue, but to date none have been addressed in the mouse model of skeletal regeneration.

**Figure 1:**
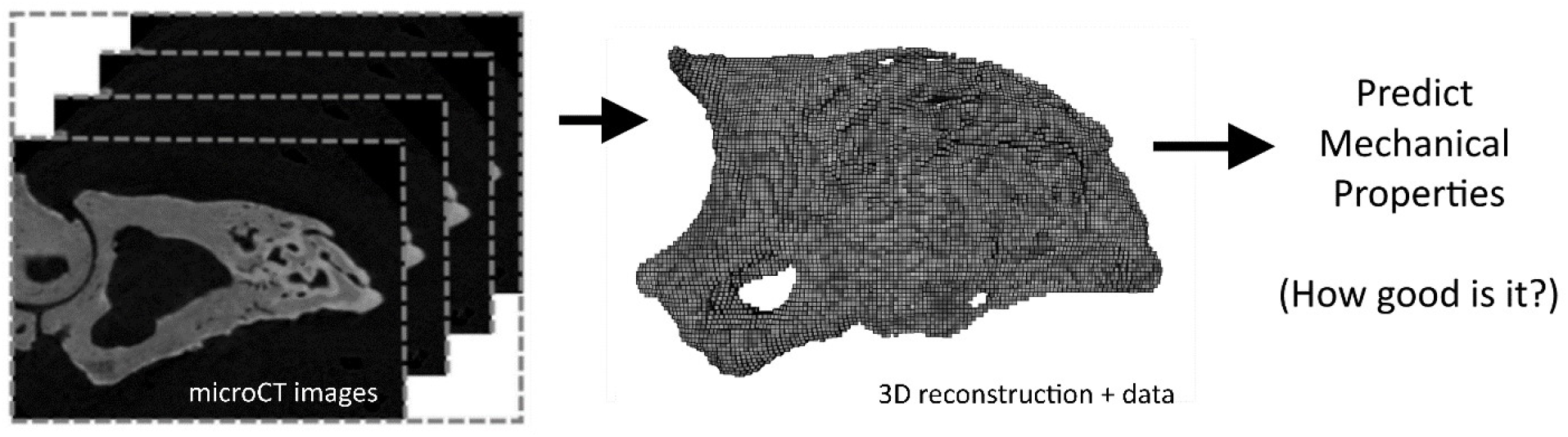
Image processing and spatial analysis on bone data from µCT image stacks offers a path to predicting mechanical properties and quality.

Nanoindentation allows experimental measurement of mechanical properties at and below the micrometer level through analysis of the recorded load-displacement curve, commonly through use of the Oliver-Pharr model (Oliver and Pharr, 2011). Nanoindentation has been widely used in other areas of bone research, including examination of stiffness versus mineralization in osteogenesis imperfecta mice, strain-dependence and viscoelastic behavior in mouse calvarial and tibial bone, the general effect of preparation and testing conditions, effect of indenter geometry, influence of embedded technique (PMMA versus epoxy), difference in spatial variation of elastic modulus within a healing callus, and heterogeneity of nanoindentation throughout the mouse tibia (Hoffler et al., 2005; Maruyama et al., 2015; Mora-Macias et al., 2017; Pepe et al., 2020; Rodriguez-Florez et al., 2013; Vanleene et al., 2012).

This research details the use of µCT images, digital image processing, and mineral density-to-elastic modulus correlation relationships to calculate the elastic modulus of regenerated and uninjured mouse P3 digits spatially throughout the whole digit, with validation through comparison to elastic modulus values calculated through nanoindentation experiments on the exact same digit samples. This allows comparative evaluation of the structural and mechanical quality of regenerated and uninjured bone.

## 2. Materials and Method

### Overview

Mouse digits were collected from young and old mice in unamputated and regenerated states, where regenerated digits were collected at 42 days past amputation (D42). These digits were then imaged through µCT scanning, followed by subsequent nanoindentation on the same digits.

### Ethics Statement

All experiments were performed in accordance with the standard operating procedures approved by the Institutional Animal Care and Use Committee of Tulane University School of Medicine.

### Amputations and Animal Handling

Adult 6-month old male and female CD1 wild type mice were purchased from Charles River (Wilmington, MA). Mice were anesthetized with 1-5% isoflurane gas with continuous inhalation. The second and fourth digits of both hind limbs were amputated at the P3 distal level as described previously (Busse et al., 2019; Sammarco et al., 2014) and regenerating digits were collected at day 42 for analysis. The sample size of mice used was N = 3, unamputated (UA) digits and N=3 regenerated (D42) digits for both numerical modeling and nanoindentation.

### Micro Computed Tomography

*Ex vivo* µCT images of mouse digits were acquired using a Bruker SkySkan 1172 scanner (Bruker, Kontich, Belgium) at 50 kV and 201 μA, with 2K resolution and an isotropic voxel size of 3.9 μm. Images were captured at a rotation angle of 0.2 with frame averaging of 5. Raw images were processed with Nrecon and DataViewer (Bruker, Kontich, Belgium). Each scan was reconstructed with a beam hardening correction of 24%, no smoothing correction, and a dynamic range of 0.00-0.339. Reconstructed output files were in 8-bit BMP format. Attenuated x-ray data values were calibrated to mineral density using standard 0.25 and 0.75 mg hydroxyapatite density phantoms and converted to greyscale output.

### Density-Elastic Modulus Calculation

Calculation of elastic modulus values utilized a processing pipeline starting with bone mineral density (BMD) as measured by µCT along with established density-elasticity relationships (Knowles et al., 2016). Calculation of elastic modulus values was performed on the entire P3 bone for unamputated digits (all cortical bone) and regenerated digits (both cortical and trabecular bone) through digital image processing and numerical calculations. Adapting recent work by the authors in semi-automated analysis of P3 digit bone mineral density (Hoffseth et al., 2021), scripts developed in Python (Rossum, 1995) code were used to process the µCT image stacks of each mouse digit and reconstruct three dimensional models. Greyscale images with calibrated pixel intensity representing mineral density (g/cm^3^) were manually cropped and segmented using FIJI to remove the second phalangeal element and any material captured in the CT image proximal to the P3 digit as part of the imaging procedure. An automated process then imported the CT images and converted image data into matrix form, with thresholding applied using a value of 60 (out of 255 greyscale, approximately 0.75 g/cm^3^) to remove out non-bone material and artifacts. Mineral density values were calculated through averaging greyscale pixel intensity using L3 sized voxels (L=3 pixels) for each representative data point though iterative operation over the digit image stack, reducing computation time without avoiding loss of digit characteristics.

Elastic modulus values were calculated from µCT measured volumetric bone mineral density values using equations relating density to elastic modulus and densitometric relationships linking density of hydroxyapatite equivalent to the ash density, an approach commonly used in CT-based finite element modeling (quantitative computed tomography, or qCT), with calibration phantoms used to calibrate BMD values (Knowles et al., 2016). Four density-modulus relationships previously used in CT-based elastic modulus calculation for bone were selected from literature (Eberle et al., 2013a, b; Edwards et al., 2013; Nishiyama et al., 2013), chosen for experimental validation in finite element studies and reporting of densitometric and scan settings. They are detailed and numbered below, and were evaluated through comparison to nanoindentation test results.

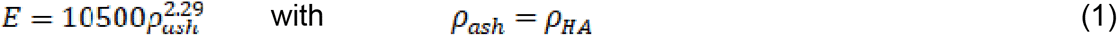

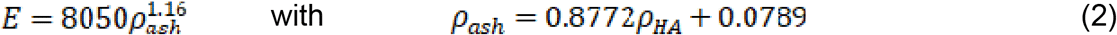

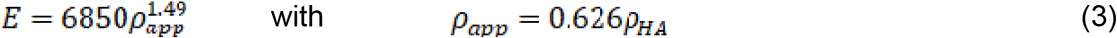

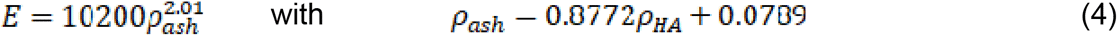

After calculation of elastic modulus values for each representative point, digits were rendered in 3D space using Python scripting and Mayavi mlab (Ramachandran and Varoquaux, 2011) to better visualize the spatial distribution of elastic modulus values throughout the magnitude range. An intensity scheme was used to enhance viewing contrast.

### Nanoindentation Measurements

Nanoindentation tests were performed by the Mayo Clinic Biomechanics Core (Rochester, MN). Mouse P3 digits were provided, stripped of all soft tissue and in frozen condition. Specimens were embedded in polymethylmethacrylate (PMMA) in acrylic cylinders. Using a combination of a low-speed diamond saw and a polishing/grinding system, the digits were sectioned along the sagittal plane. Once a bone’s cross section was revealed, it was manually polished using successively finer abrasive cloths (400, 600, 800, and 1200 grit) with a final polish using a micro cloth and slurry of 0.05-μm aluminum abrasive. Indentation testing was conducted on the cortical bone with a nano-indentation system (TI 950, Hysitron, Minneapolis, MN) equipped with a diamond Berkovitch pyramidal tip. A total of 8 sites, 5 distributed across the distal area and 3 distributed across the proximal area, were tested on each bone. At each site, a 2 × 2 array was indented with 15-μm spacing between indents. Indentation was conducted under load control at a rate of 500 μN/s to a peak load of 2,000 μN with a 60-s hold before unloading to reduce viscoelastic effects. The reduced modulus (Er; GPa), and hardness (H; GPa), were calculated from load versus displacement data using the Oliver–Pharr model (Oliver and Pharr, 2011), with resulting elastic modulus (E; GPa) calculated from the reduced modulus using Poisson’s ratio = 0.3 for bone and E=1141 GPa, Poisson’s ratio = 0.07 for the diamond indenter.

### Statistics

Elastic modulus (E) and hardness (H) were analyzed using two-way ANOVA models for bone area (proximal or distal) and amputation status (UA or D42) with random effects at the level of mouse and nanoindentation site within mouse using the R package nlme (Pinheiro, 2021). Statistical significance was assessed by testing the set of relevant contrasts (UA proximal vs distal, D42 proximal vs distal, and UA distal vs D42 distal) with adjustment for multiple comparisons using the R package multcomp (Hothorn et al., 2008). Numerically calculated elastic modulus values were reduced to 1000 values per digit and location (proximal or distal) via random sampling. Samples were analyzed with two-way ANOVA models for bone area and amputation status with random effects at the mouse level using nlme and multcomp as described above. Local polynomial regression curves were fit to the full set of calculated elastic modulus values for each digit as a function of proximal-distal location using the R function “loess” with span parameter set to 0.33.

## 3. Results

Values for E were determined through numerical calculation and experimental measurement for all 6 digits, with subdivision into two categories, D42 and UA. Experimental measurement of E was performed using nanoindentation, with 32 measurements total for each digit in regionally specific locations (proximal and distal) to validate and compare with numerical values. H values were obtained from the same measurements. The nanoindentation-measured E and H values were compared between the proximal and distal areas of the bone and between the unamputated and regenerated sample groups. E values may be seen in both Figure 2 and Table 1. H values may be found in the Appendix. Measured E and H values showed no significant difference between the proximal and distal areas of the unamputated digit, however in a regenerated digit (D42) the E values are lower in the distal regenerated bone than in the proximal uninjured stump (p<0.001, 95% CI for difference = 0.93, 3.78), and similar for H (p<0.001, 95% CI for difference = (0.05, 0.18). A comparison between groups shows that the E values for the distal region in regenerated bone are also significantly lower than those in an unamputated digit (p<0.001, 95% CI for difference = -2.95, -1.02), with similar trend for H (p< 0.001, 95% CI for difference = (−0.15, -0.06).

**Figure 2.**
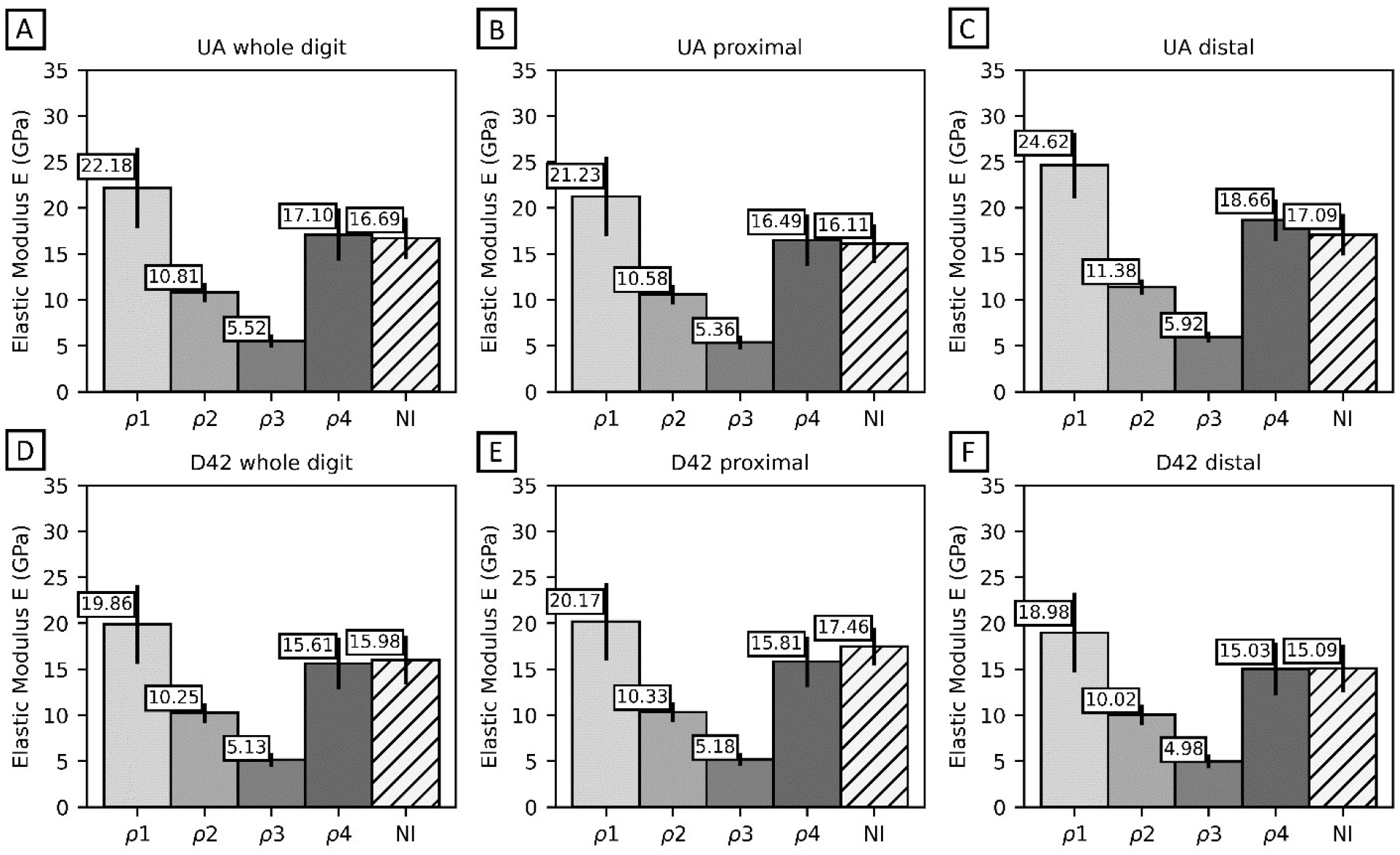
Comparison of calculated elastic modulus values from all four µCT density-modulus relationships (noted by ρ1, ρ2, ρ3, ρ4, respectively) with those measured by nanoindentation experiment (noted by NI), for a) whole UA digits, b) UA proximal regions, c) UA distal regions, d) whole D42 digits, e) D42 proximal regions, and f) D42 distal regions. Relationship 4 shows best overall agreement with experimentally measured values.

**Table 1:** Elastic Modulus Values from µCT calculations, and nano indentation experiments, for whole digits mean values, with standard deviation in parenthesis.

| Sample | Elastic Modulus, GPa |  |
| --- | --- | --- |
|  | Calculated | Experiment |
| D42 233 | 15.789 (2.799) | 16.612 (3.103) |
| D42 234 | 16.067 (2.915) | 15.496 (2.559) |
| D42 235 | 14.959 (2.647) | 15.829 (2.235) |
| UA 233 | 16.634 (2.738) | 16.453 (2.242) |
| UA 234 | 17.447 (3.046) | 15.263 (1.994) |
| UA 235 | 17.224 (2.718) | 18.354 (2.476) |

Numerical calculation of E used µCT measured values of bone mineral density to calculate predicted E values for every representative voxel data point in the reconstructed digit. These values were then compared against experimental values. Several different density-modulus relationships were evaluated in calculation of E values, where results of the evaluation may be seen in Figure 2, showing comparison of the calculated E value using select relationships detailed in Equations 1-4 versus the measured E value from nanoindentation. Comparing results shows Equation (4), with the associated hydroxyapatite-to-ash density relationship, returned values which showed best agreement with values measured through nanoindentation. Calculated average E values for whole digits are displayed in Table 1, compared to each average E value obtained from experimental measurement. Relation 4 showed the best agreement across all the unamputated and regenerated digit groups, with an average error of 5.3% across all four UA and D42 distal and proximal regions, and was used for further analysis.

The numerically calculated E values from both digit groups were able to statistically pick up spatial changes in elasticity that analysis on the experimental E values did not detect. Analysis identified an increase in the E values in the distal end of the unamputated digit (p<0.001, 95% CI for proximal-distal difference = (−3.52, -3.27), as well as confirmed decreases in the distal end of the D42 regenerated bone as compared to both the proximal stump (p<0.001, 95% CI for difference = (0.04, 1.20), and the distal unamputated tip (p<0.001, 95% CI for difference = (5.64, 5.90). Additional results may be seen in Figures 3 and 4 investigating trends from proximal to distal ends of each digit.

**Figure 3.**
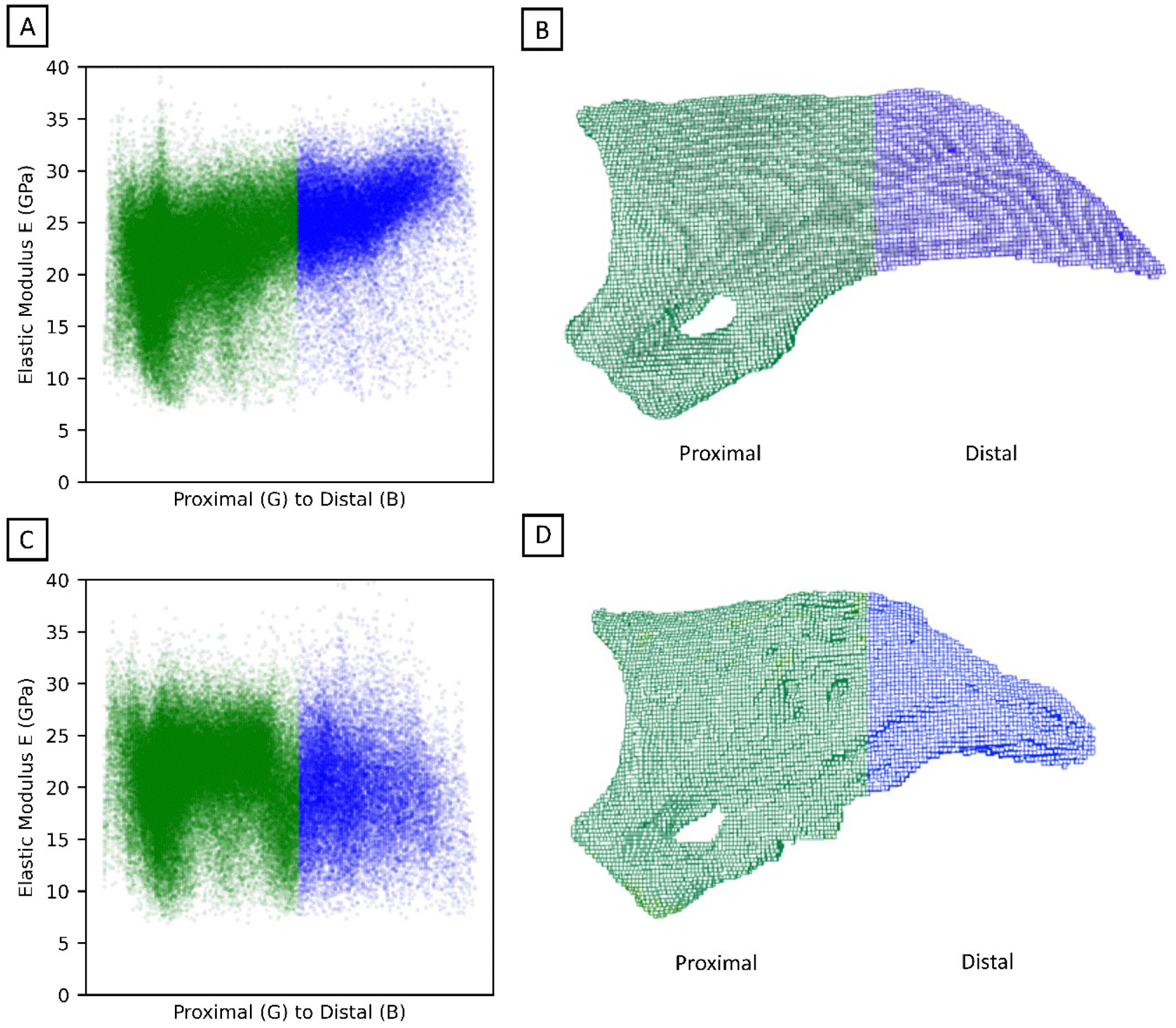
Example Proximal to Distal E distribution, for representative regenerated and unamputated digits. Green and blue colors distinguish proximal and distal regions. A) scatterplot where the Y-axis is Elastic Modulus, and X-axis is distance from center plane in voxels, showing the distribution of modulus magnitudes from proximal to distal ends. B) Matching rendering of the actual digit, displaying the proximal and distal spatial regions.

**Figure 4.**
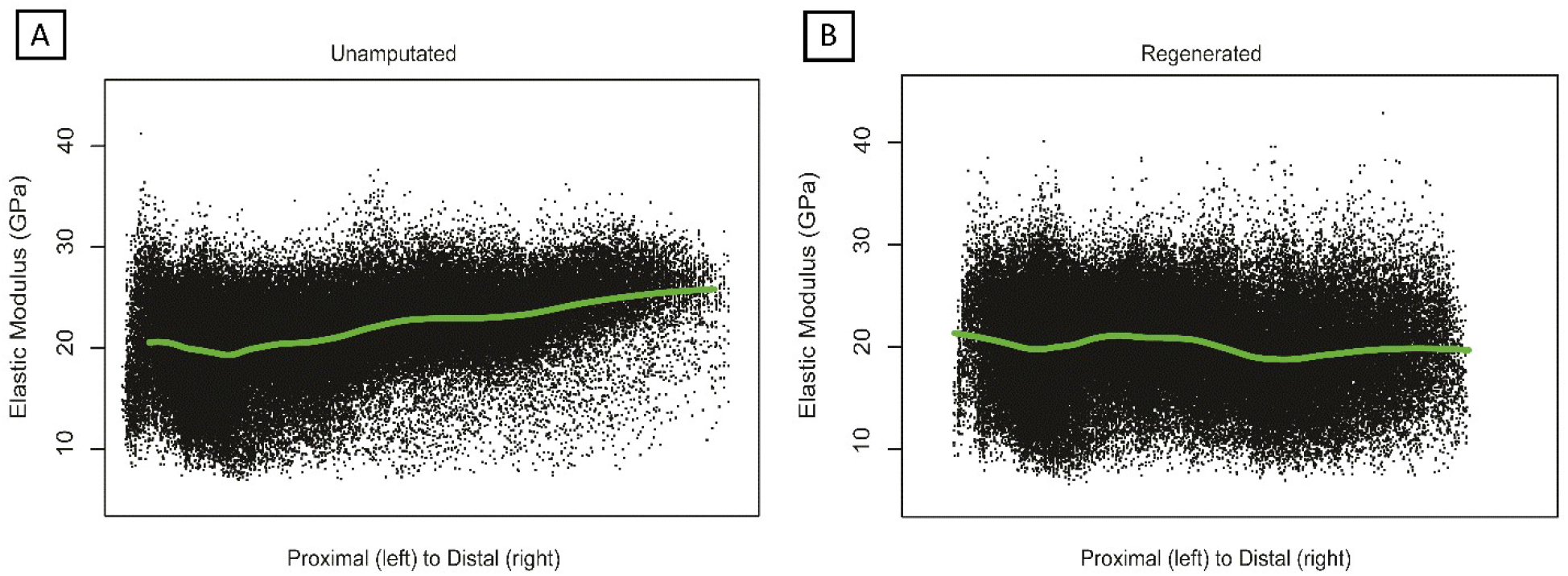
Plotting of numerically calculated Unamputated (A) and Regenerated (B) elastic modulus values versus proximal to distal distance. Local polynomial regression curve indicated in green, indicating the trend in modulus value from proximal to distal.

Spatial representation of E values in the proximal and distal regions of UA and D42 digits can be seen in Figure 3. Figure 3a displays results of plotting individual voxel E values versus their distance from the plane dividing proximal and distal regions for a representative example UA digit. Figure 3b shows the 3D render of the UA digit from µCT images, with respective proximal and distal regions. Figures 3c and 3d show the same for a D42 digit. Comparing Figure 3a and Figure 3b, the rising trend of the blue distal region in the UA digit is different than the less tightly packed distribution of values of the D42 distal region, where there appears to be a neutral or slight downward trend in E values. Figure 4 shows the E magnitude distribution for all digits in each group, with calculated E versus the distance from proximal to distal. A local polynomial regression curve demonstrates the variation in trend of E from the proximal to distal end of the digit.

Calculation of the E at every spatial point in the rendered 3D digit volume is seen in Figure 5, for representative examples of both UA and D42 digits (Figures 5a and 5c, respectively). The spatial distribution of E values visually appears to be more uniform in unamputated digits, than in regenerated digits. Additionally, the proximal end comprising the original cortical bone is predictably more uniform in than the distal region in D42 digits. Figures 5b and 5d display a midplane cut of the digit for UA and D42, respectively, showing the inner distribution of elastic values, where the disorganization and increased internal void network is clearly seen in the D42 digit.

**Figure 5:**
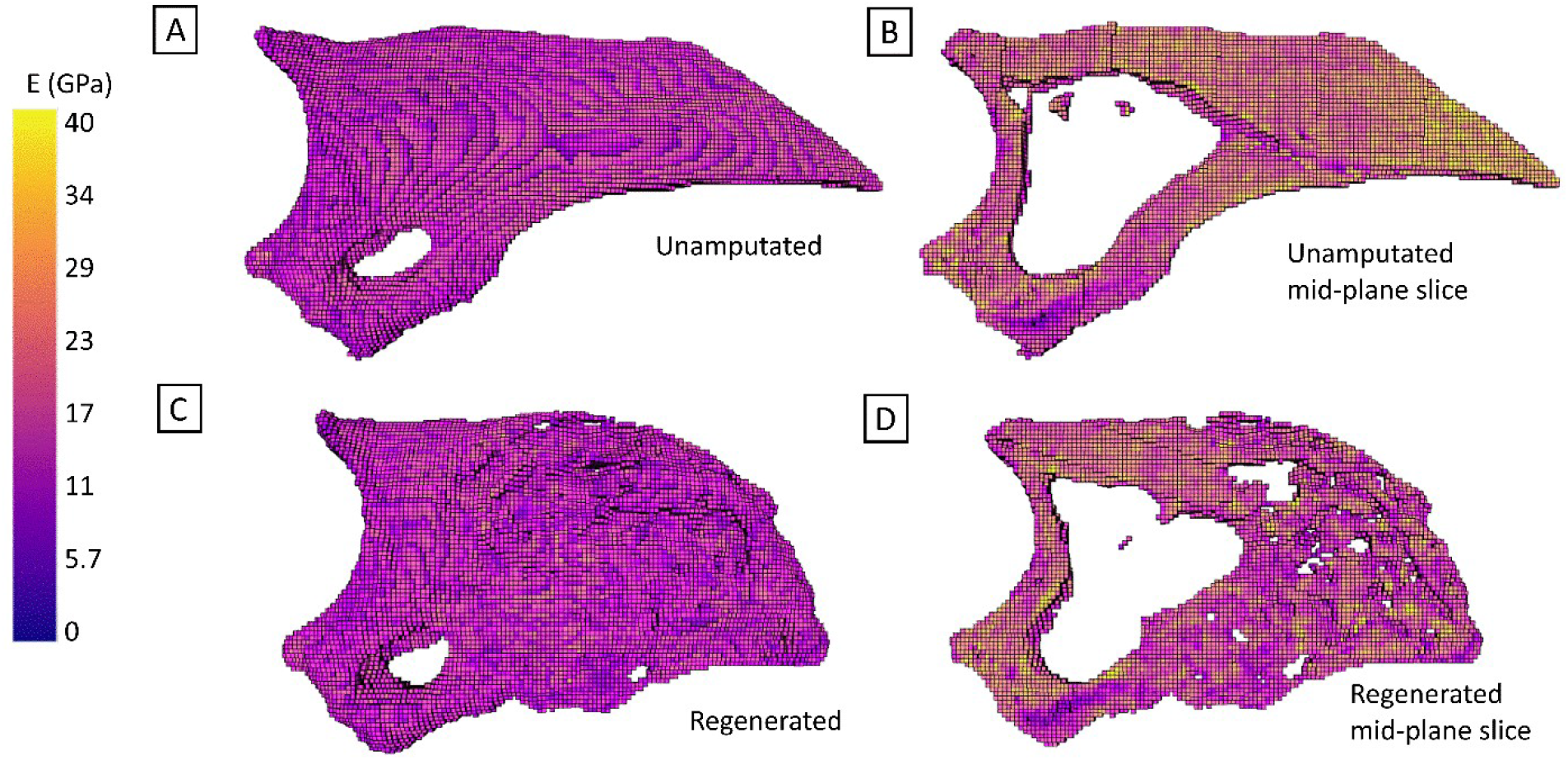
Intensity Depiction in 3D of Elastic Modulus (E) values, calculated from mineral density data measured through µCT image stack analysis, with A) whole digit rendering of Digit 150 D42, and B) a planar vertical slice along the proximal to distal axis of the same digit, showing internal spatial distribution of the elastic modulus values.

## 4. Discussion

Improved, quantifiable understanding of regenerated bone quality in terms of mechanical properties is critical for developing and confirming improved bone health treatments for both young and aged populations. Analyzing µCT data with traditional bone morphometric approaches to quantify and visualize bone architecture has proven a valuable, but unable to fully characterize structurally relevant, mechanical properties on their own. The research presented here combines semi-automatic methods of image processing and densiometric-based numerical calculation of elastic values to predict spatially distinct elasticity values in regenerated bone, and builds a bridge between analysis of µCT data and measured mechanical response through validation by nanoindentation measurements. Numerical calculation of elastic values not only improves analysis of the structural quality of regenerated bone in the mouse digit model, but provides a predictive method with statistically relevant sensitivity that was not possible with scale-associated limitations in mechanical testing.

Collectively, these results highlight the importance of using predictive modeling to assess the biomechanics of regenerated bone, and applicability of common densiometric relationships used in qCT studies. Furthermore, using nanoindentation to more rigorously assess the elasticity of the bone strengthens this approach and helps surmount some impediments to investigating biomechanics in bones restricted by shape and size, such as challenges and limitations in the mouse digit regeneration model. The difficulties inherent in very small bone sample mechanical testing can be large, where effective spatial measurement is prohibitive due to time, cost, or sample fragility. However, image-based numerical approach detailed here overcomes these impediments and provides a greater breadth of analysis for evaluating mechanical properties in the mouse digit model.

Subsequently, improved analysis allows a more focused look at how bone tissue rebuilds spatially, proximal to distal, after amputation. Numerically calculated and experimentally measured values of E show agreement both for regenerated and unamputated digits. Comparing calculated results for distal and proximal regions shows that UA digits appear to have a higher E near the distal tip as opposed to the proximal end near the joint, where D42 regenerated bone does not appear to have the same trend, instead showing a decline in E values. Similarly, experimental hardness values exhibit a similar trend. Together these data indicate that newly regenerated bone is less able to resist permanent deformation and maintain shape, and is curious in light of the fact that we recently showed an increase in mineral density values in newly regenerated bone (Hoffseth et al., 2021) (evident in these analysis as the higher individual distal values). This may be explained by a difference in bone tissue organization, where woven bone is generally more dense than lamellar bone, but lamellar bone if oriented in direction parallel to loading may perform just as well or better in terms of displacement and resistance to loading, due to its anisotropy (Currey, 2002). This underscores the continued need to evaluate the mechanics and structure of regenerated bone to better assess potential treatments both in the research phase and in clinic.

A benefit of processing of spatial 3D data from µCT images is visualization and discovery of more changes in rebuilding bone. Evaluation of the proximal to distal distribution of E values in the digits show that the values are overall more broadly distributed in magnitude in the rebuilt distal region than in the uninjured distal region. This may be interpreted as a more disorganized construction than in neonatal development.

Utilization of µCT based calculation of E values validated through nanoindentation experiments to analyze regenerated mouse digit bone is a step forward in assessing the mechanistic, structural quality of regenerated bone. The mouse digit regeneration model was chosen for its value in determining differences between regenerated and non-regenerated bone, and where its unique challenges had been a barrier to more detailed structural analysis. Following the path from qCT to finite element analysis (qCT-FE) in the future holds potential for further analysis in subsequent work, now that the link between µCT based calculation and experimental assessment of E has been established in the mouse digit model of skeletal regeneration.

## 5. Conclusion

This work demonstrates and validates the use of density-modulus relations in calculation of elastic modulus for regenerated mouse P3 bone based on micro-computed tomography (µCT) images through comparison to nanoindentation experiments, and thus allows for improved evaluation and analysis of structural quality when assessing bone regrowth success. Evaluation of select density modulus relationships showed a common variation of the power law density-modulus relation used in qCT-FE methods returned best agreement with experimental results. Spatial rendering of µCT originating calculated modulus values throughout the 3D digit allows more powerful analysis of the actual bone structure being rebuilt. Application to regenerated and uninjured mouse bone shows new insight into the regeneration process, with regenerated bone displaying lower elastic and hardness values than uninjured.

## Acknowledgements

Funding was provided by a research grant from the National Institute of General Medical Sciences P20GM103629 (Sammarco), and research start-up funds (Hoffseth) from Louisiana State University.

## Appendix

**Table A1:**
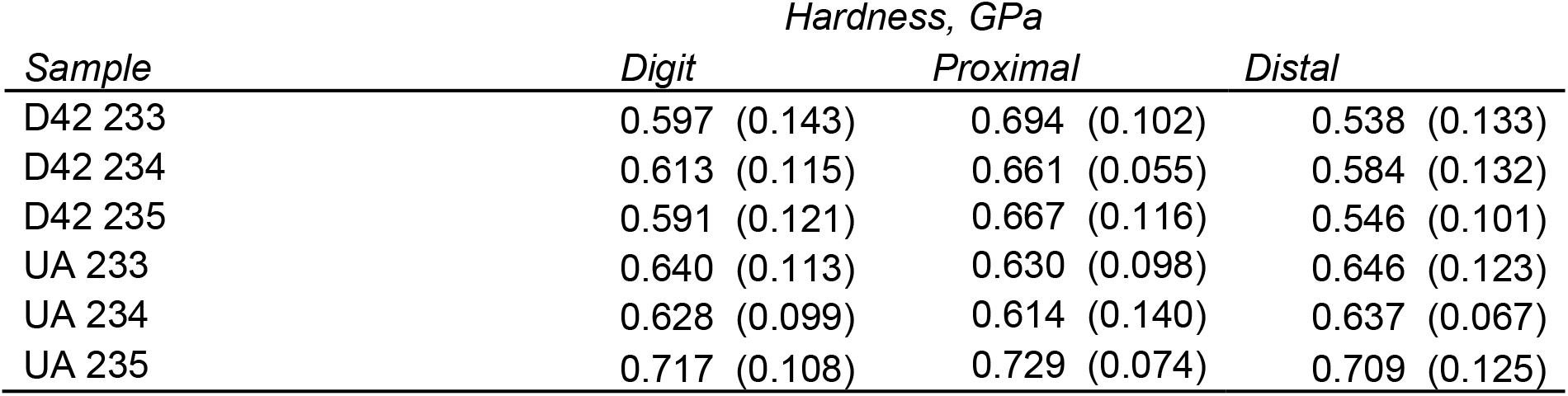
Hardness values (GPa) measured by nanoindentation experiment. Values for overall digit, proximal region only, and distal region only, with standard deviation in parentheses.

